# Functional divergence of the *bag of marbles* gene in the *Drosophila melanogaster* species group

**DOI:** 10.1101/2021.06.25.449946

**Authors:** Jaclyn E. Bubnell, Cynthia K.S. Ulbing, Paula Fernandez Begne, Charles F. Aquadro

## Abstract

In *Drosophila melanogaster*, a key germline stem cell (GSC) differentiation factor, *bag of marbles* (*bam*) shows rapid bursts of amino acid fixations between sibling species *D. melanogaster* and *D. simulans*, but not in the outgroup species *D. ananassae*. Here, we test the null hypothesis that *bam’*s differentiation function is conserved between *D. melanogaster* and four additional *Drosophila* species in the *melanogaster* species group spanning approximately 30 million years of divergence. Surprisingly, we demonstrate that *bam* is not necessary for oogenesis or spermatogenesis in *D. teissieri* nor is *bam* necessary for spermatogenesis in *D. ananassae*. Remarkably *bam* function may change on a relatively short time scale. We further report tests of neutral sequence evolution at *bam* in additional species of *Drosophila* and find a positive, but not perfect, correlation between evidence for positive selection at *bam* and its essential role in GSC regulation and fertility for both males and females. Further characterization of *bam* function in more divergent lineages will be necessary to distinguish between *bam’s* critical gametogenesis role being newly derived in *D. melanogaster, D. simulans, D. yakuba*, and *D. ananassae* females or it being basal to the genus and subsequently lost in numerous lineages.

## Introduction

In most sexually reproducing animals, egg and sperm production begins with the differentiation of germline stem cells. These germline stem cells (GSCs) divide both to self-renew to maintain the germline and to produce daughter cells that differentiate to produce gametes (Kahney et al. 2019). The cellular processes of daughter cell differentiation and stem cell self-renewal are the most fundamental functions of stem cells and are therefore highly regulated and presumed to be conserved. In the germline, the mis-regulation of either of these processes leads to sterility (Gleason et al. 2018). However, many of the genes that control GSC maintenance and GSC daughter differentiation in *D. melanogaster* are rapidly evolving due to positive selection for amino acid diversification (Civetta et al. 2006; Bauer DuMont et al. 2007; Choi and Aquadro 2015; Flores, DuMont, et al. 2015).

One such gene is *bag-of-marbles* (*bam*) which exhibits a rapid burst of amino acid substitutions between the sister species *D. melanogaster* and *D. simulans*, acts as the key switch for GSC daughter cell differentiation in *D. melanogaster* females, and is necessary for terminal differentiation of spermatogonia in males (McKearin and Spradling 1990; Civetta et al. 2006; Bauer DuMont et al. 2007; Insco et al. 2009). Despite 60 fixed amino acid differences between *D. melanogaster* and *D. simulans*, a *D. simulans bam* transgene is sufficient for GSC daughter cell differentiation in *D. melanogaster* females and males. The intriguing observations that *bam* is both rapidly evolving due to positive selection and functions as a switch for GSC daughter differentiation in *D. melanogaster* has led us to investigate what could be driving its positive selection between species (Flores, Bubnell, et al. 2015).

In females, *bam* is repressed in GSCs, and upon its expression promotes differentiation by binding to the protein product of *benign gonial cell neoplasm* (*bgcn)* as well as a series of other protein partners to repress the translation of self-renewal factors *nanos* and *eIF4a* (Li et al. 2009; Ting 2013). The GSC daughter cell differentiates into a cystoblast, which then goes through four rounds of mitotic divisions while *bam* localizes to the fusome, a cellular structure that interconnects the resulting dividing cells that make up the cyst. In these cells, Bam functions with Bgcn to regulate mitotic synchrony (Chen and McKearin 2003). Before meiosis begins, *bam* expression is repressed. In males, *bam* is expressed in GSCs and its expression increases as differentiation progresses (Insco et al. 2009). In early dividing spermatogonia, Bam protein must reach a threshold to bind with partners Bgcn and *tumorous testis* (*tut*) to repress *mei-P26* and switch the cellular program from proliferation to terminal differentiation to begin meiosis (Chen et al. 2014). After this switch, *bam* expression is rapidly repressed. Loss of *bam* function in either sex results in the over-proliferation of either GSCs in females or spermatogonia in males causing tumors and sterility (McKearin and Spradling 1990).

One possible explanation for the observed signal of positive selection at *bam* is a change of function. As *bam* regulates gametogenesis, it is possible that life history or environmental factors drive positive selection at *bam*. It is even possible that the genetic interaction between the endosymbiotic bacteria *Wolbachia pipientis* and *bam* has led to such changes in *bam*’s function in different lineages, as *Wolbachia* infects the GSC niche and is propagated through the germline (Flores, Bubnell, et al. 2015). While functional divergence of *bam* seems less likely due to its essential role in GSC maintenance and differentiation in *D. melanogaster, bam* sequence is highly divergent among species. It is even difficult to confidently align *bam* sequence between *D. melanogaster* and *D. ananassae*, both members of the *melanogaster* species group (subgenus Sophophora), let alone the many more divergent *Drosophila* species (Bauer DuMont et al. 2007). Additionally, *bam* appears to be a novel gene in the genus *Drosophila* as no orthologues are reported in any non-*Drosophila* genomes using orthoDB v10 (Kriventseva et al. 2019) and Ensembl Metazoa (Howe et al. 2020) (both accessed 11/4/2021).

Here, we investigate the possibility that the observed signals of positive selection in *D. melanogaster* and *D. simulans* at *bam* are due to selection for a novel function for *bam* in gametogenesis in these lineages. To do this, we ask if *bam* is necessary for gametogenesis in diverse species within the *Drosophila* genus. We generate *bam* loss-of-function alleles in five *Drosophila* species in the *melanogaster* species group (*D. melanogaster, D. simulans, D. yakuba, D. teissieri*, and *D. ananassae*). Surprisingly, we find that *bam* is not necessary for gametogenesis in one or both sexes in two species within the *melanogaster* species group. We also analyze additional population samples of species mainly within the *melanogaster* species group and a few more distant lineages for nucleotide sequence variation at *bam*. Within the *melanogaster* species subgroup (*D. melanogaster, D. simulans, D. yakuba, D. santomea*, and *D. teissieri*), we find evidence of lineage-specific signals of adaptation for three species, but not for *D. teissieri* or *D. santomea*. Outside of the *melanogaster* species subgroup, we find similar patterns of lineage-specific signals of positive selection. Together with our *bam* loss-of-function results, these observations raise the possibility that the critical role for *bam* in GSC differentiation is newly derived in the *melanogaster* species group and the signatures of positive selection we observe in some of those species is associated with species-specific refinements of *bam*’s GSC differentiation function or species-specific adaptive pressures acting on *bam*’s GSC differentiation function.

## Results

### Generating *bam* null alleles for non-*melanogaster* species

To ask if *bam’s* GSC function is conserved in species outside of *D. melanogaster*, we designed and generated a series of *bam* null alleles in five species in the *melanogaster* species group to assay for GSC daughter differentiation (Fig 1A). To do this, we used CRISPR/Cas9 to introduce a 3xP3-YFP or a 3xP3-DsRed gene cassette into the first exon of *bam*, thereby disrupting the *bam* coding sequence and introducing a premature termination codon (Fig 1B). This also generates an allele that is trackable by eye color, which is necessary for the non-*melanogaster* species of which we do not have balancer chromosomes to maintain alleles that cause sterility. To generate the homozygous *bam* null genotype, we cross the 3xP3-YFP line to the 3xP3-DsRed line and select flies with both DsRed and YFP positive eyes. This scheme also allows us to use the same cross to assay the heterozygous and wildtype *bam* siblings (Table 1).

**Fig 1.**
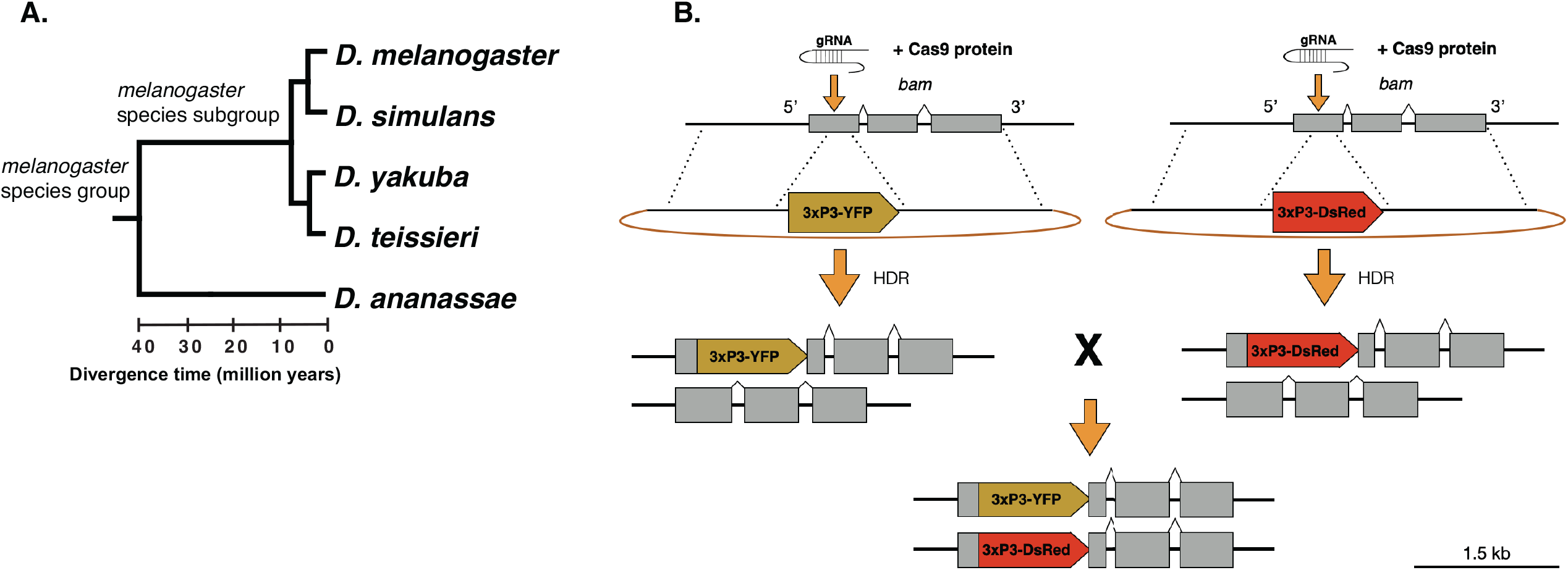
Species chosen to generate *bam* null alleles using CRISPR/Cas9 to disrupt *bam* with a trackable eye marker cassette. (A) The five species we chose to generate *bam* null alleles span approximately 30 million years of divergence. (B) Schematic for generating null alleles with trackable eye markers using CRISPR/Cas9. We generate two null alleles using different color eye markers and cross the heterozygotes together to generate an easily trackable null genotype.

**Fig 2.**
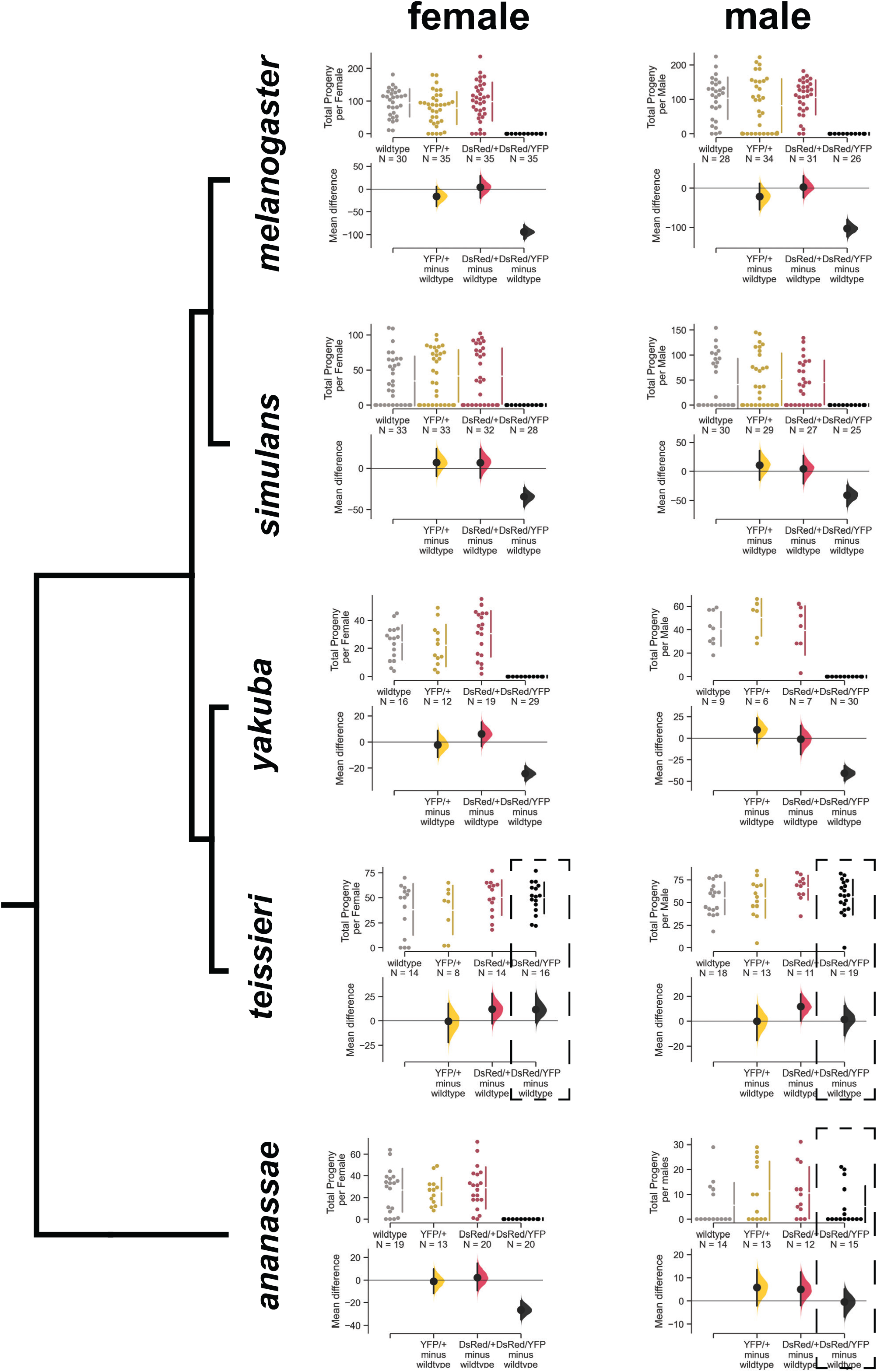
*Bam* is necessary for female and male fertility in *D. melanogaster, D. simulans, and D. yakuba*, but not for *D. ananassae* males nor both *D. teissieri* males and females. Swarm plots showing the total progeny per female or per male for each *bam* genotype and Cumming estimation plots showing the 95% confidence interval effect size. Dotted boxes show genotypes where the phenotype did not match that of *D. melanogaster*. For *D. melanogaster, D. simulans, and D. yakuba*, the *bam* null DsRed/YFP females and males were sterile (P < 0.0001, permutation test), and the heterozygous genotypes did not have a significant effect on fertility (P > 0.05, permutation test). For *D. teissieri*, the *bam* null DsRed/YFP females and males did not have significantly different fertility from wildtype, nor did the heterozygotes. For *D. ananassae*, the *bam* null DsRed/YFP females were sterile (P < 0.0001, permutation test), and the heterozygotes did not affect fertility. However, the *bam* null DsRed/YFP males were fertile and there was no significant difference between the null and heterozygotes and wildtype fertility (P>0.05, permutation test).

**Table 1.**
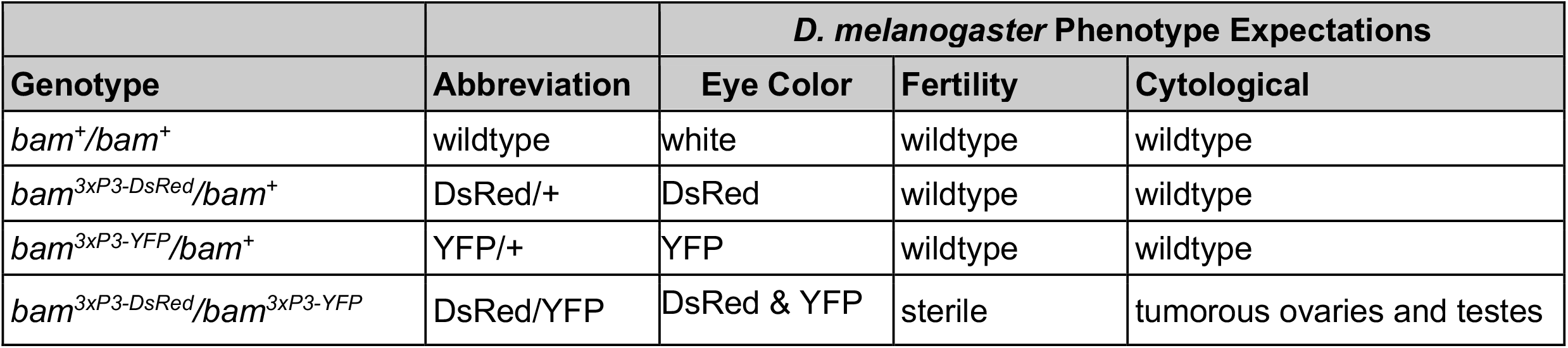
Abbreviations for *bam* genotypes for all species and the expected phenotypes as observed in *D. melanogaster*.

| Genotype | Abbreviation | *D. melanogaster* Phenotype Expectations |  |  |
| --- | --- | --- | --- | --- |
|  |  | Eye Color | Fertility | Cytological |
| *bam*<sup>+</sup>/*bam*<sup>+</sup> | wildtype | white | wildtype | wildtype |
| *bam*<sup>3xP3-DsRed</sup>/*bam*<sup>+</sup> | DsRed/+ | DsRed | wildtype | wildtype |
| *bam*<sup>3xP3-YFP</sup>/*bam*<sup>+</sup> | YFP/+ | YFP | wildtype | wildtype |
| *bam*<sup>3xP3-DsRed</sup>/*bam*<sup>3xP3-YFP</sup> | DsRed/YFP | DsRed & YFP | sterile | tumorous ovaries and testes |

We used *D. melanogaster* as a control to ensure our method would generate a loss of function *bam* phenotype, as the *bam* null phenotype has been well characterized in this species (Table 1). We also generated *bam* nulls in four other species within the *melanogaster* species group *D. simulans, D. yakuba, D. teissieri*, and *D. ananassae* (Fig 1A).

### *bam* is necessary for female and male fertility in *D. melanogaster, D. simulans*, and *D. yakuba*, but only in *D. ananassae* females, and in neither *D. teissieri* males nor females

In *D. melanogaster*, both male and female *bam* nulls are sterile as early germline cells fail to differentiate and proceed through gametogenesis. In classic null alleles of *D. melanogaster bam* (*bam*^*Δ59*^ and *bam*^*Δ86*^), one copy of functional *bam* is sufficient to rescue the *bam* null sterility phenotype. We first asked if our novel *bam* null alleles behaved this way in *D. melanogaster*. Heterozygotes of each 3xP3-DsRed or 3xP3-YFP disruption line showed similar fertility to wildtype *bam* in both males and females (mean difference of adult progeny, not significant, permutation test) and transheterozygous males and females carrying both the 3xP3-DsRed and the 3xP3-YFP disruptions (which we will term the *bam* null genotype) were completely sterile (mean difference of adult progeny, P < 0.0001, permutation test) (Fig 1, File S2). We observed this same pattern for *D. simulans* and *D. yakuba* (Fig 1, File S2). We observed some sterile *D. simulans* females and males across all *bam* genotypes, which we attribute to the strain and unlikely to be related to *bam* (Fig 1, File S2). In contrast, in *D. teissieri* we found that there was no difference in fertility between any of the *bam* genotypes tested for females or males (mean difference of adult progeny, not significant, permutation test) (Fig 1, File S2). The *bam* null genotype did not result in sterility, or even reduced fertility. In *D. ananassae, bam* null females were sterile (mean difference of adult progeny, P < 0.0001, permutation test), and heterozygotes of each 3xP3-DsRed or 3xP3-YFP disruption did not significantly reduce fertility (mean difference of adult progeny, not significant, permutation test). However, in *D. ananassae* males, *bam* nulls were fertile, with no significant mean difference in progeny across the tested *bam* genotypes (Fig 1, File S2). Therefore, *bam* is not necessary for fertility in all species in the *melanogaster* species group, or even all species within the *melanogaster* species subgroup.

### Consistent with the fertility results, *bam* is also necessary for germ cell differentiation in *D. simulans* and *D. yakuba* females and males, but not in *D. teissieri* males or females, and only *D. ananassae* females

The ovaries of *D. melanogaster bam* null females feature over-proliferating germline stem cells that fail to differentiate into cystoblasts (McKearin and Spradling 1990; Ting 2013). This results in the germarium and developing downstream cysts accumulating small, GSC-like cells resembling tumors (Fig 3A). In the testes of *D. melanogaster bam* null males spermatogonia differentiation is blocked. The resulting testes feature over-proliferating spermatogonia that fail to develop into terminally differentiated spermatocytes (Fig 3A) (Insco et al. 2009; Insco et al. 2012).

**Fig 3.**
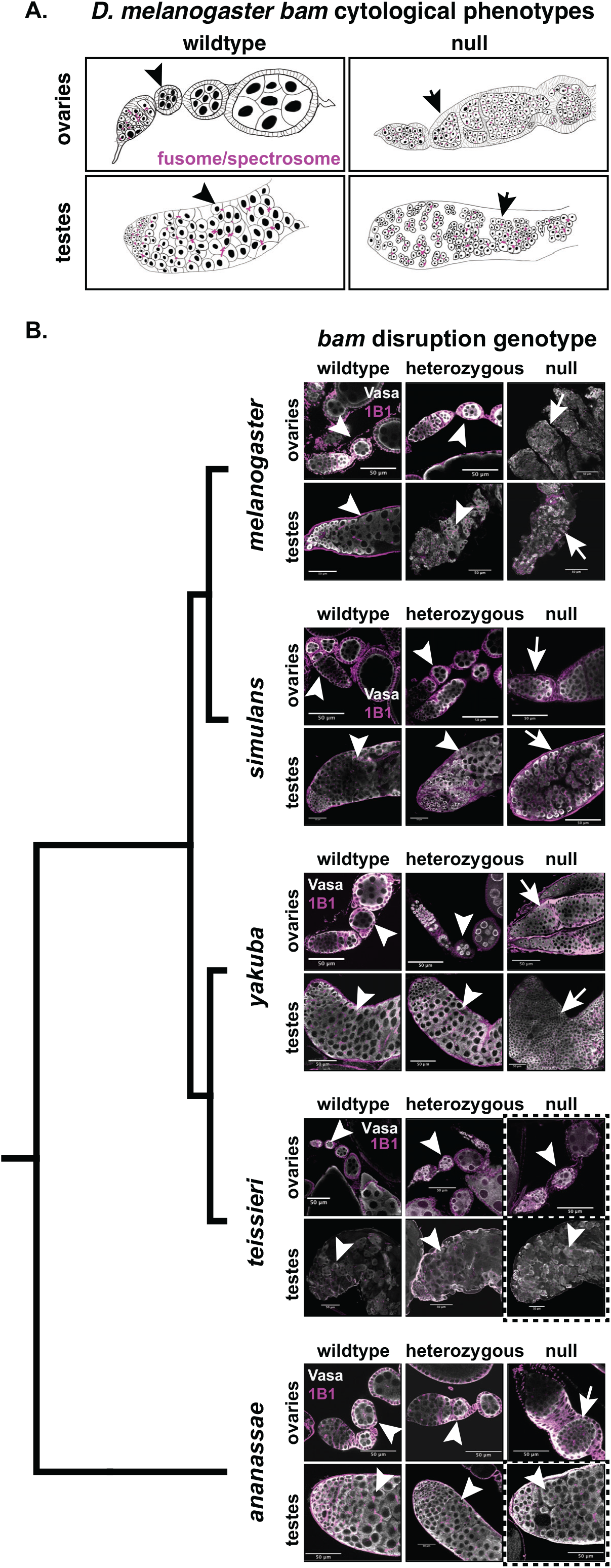
*Bam* is necessary for female and male germline stem cell differentiation in*D. melanogaster, D. simulans, and D. yakuba*, but not for *D. ananassae* males nor *D. teissieri* males and females. (A) Schematics of ovaries and testes showing the *bam* wildtype and null cytological phenotypes in *D. melanogaster*. Wildtype *bam* ovaries exhibit developing egg chambers with differentiated nurse cells and testes exhibit large, differentiated spermatocytes. Null *bam* ovaries exhibit egg chambers filled with small GSC-like tumors that contain undeveloped fusomes and testes exhibit cysts filled with small over-proliferating spermatogonia. (B) Ovaries and testes from all five species of wildtype (*bam*^*+*^/*bam*^*+*^), heterozygous (YFP*/+* or DsRed/+), and null (DsRed/YFP) *bam* genotypes. All tissue is labeled with antibodies to Vasa (germline) and Hts-1B1 (fusome). Dashed boxes indicate genotypes and tissue that are not consistent with the *D. melanogaster* phenotype. Ovaries and testes from all five species exhibit developing nurse cells or spermatocytes (respectively) for wildtype and heterozygous *bam* genotypes. We did not observe any tumorous cysts (arrowheads). Ovaries and testes from *bam* null *D. melanogaster, D. simulans, and D. yakuba* all exhibited cysts filled with GSC-like tumors (ovaries) or over-proliferating spermatogonia (testes) (arrows). The *bam* null ovaries exhibit undeveloped fusomes. *D. ananassae bam* null ovaries exhibited the tumorous egg chamber phenotype (arrow), however testes exhibited large developing spermatocytes (arrowhead) and we did not observe any tumors. The *bam* null *D. teissieri* ovaries and testes exhibited nurse cell positive egg chambers and developing spermatocytes (respectively) (arrowheads) and we did not observe any tumors.

We imaged ovaries from 2-5 day old *D. melanogaster* females from each *bam* genotype. We assayed for the characteristic *bam* null tumor phenotype using antibodies to Vasa to visualize the morphology of the germline and Hts-1B1 to visualize the fusome, which fails to fully form in *bam* mutants (Lavoie et al. 1999; Hinnant et al. 2020). We found that the *bam* null genotype resulted in the characteristic *bag of marbles* phenotype consistent with classic *bam* null alleles, with tumorous ovaries filled with undifferentiated GSC-like cells (Fig 3B). One copy of wildtype *bam* was sufficient to rescue the *bam* null phenotype, as has been previously observed for other *bam* null alleles (Fig 3B) (Lavoie et al. 1999; Flores, Bubnell, et al. 2015). Therefore, our 3xP3-DsRed and 3xP3-YFP disruptions successfully recapitulate the *bam* null phenotype in females. We imaged testes from 3-5 day old *D. melanogaster* males and also immunostained with antibodies to Vasa to label the germline and Hts-1B1 to label the fusome. We found that the *bam* null genotype resulted in tumorous testes filled with small over-proliferating cells that fail to differentiate into spermatocytes (Fig 3B). We found that one copy of wildtype *bam* was sufficient to rescue the *bam* null tumor phenotype, as has been previously reported for classic *bam* null alleles (Gönczy et al. 1997; Shivdasani and Ingham 2003)(Fig 3B). Therefore, our *bam* null alleles also recapitulate the previously described *bam* null phenotype in *D. melanogaster* males.

We repeated the same analysis for the other species in the study. We found for *D. simulans* and *D. yakuba* both female and male homozygous *bam* null genotypes exhibited the characteristic ovarian and testes tumors as described in *D. melanogaster* (Fig 3B). In these two species for both males and females, one copy of *bam* was also sufficient to rescue the *bam* null phenotype as observed in *D. melanogaster* (Fig 3B). However, in the *D. teissieri bam* null genotype, we observed no evidence of GSC over-proliferation in females or spermatogonia over-proliferation in males. We observe the same phenotype for the heterozygous alleles in both males and females, as well as wildtype (Fig 3B). This indicates that *bam* is not necessary for GSC daughter differentiation in *D. teissieri* females nor is *bam* necessary for spermatogonia differentiation in males. In *D. ananassae bam* null females, we observed the characteristic *bam* null phenotype observed in *D. melanogaster* with over-proliferating GSC-like cells in the germarium and cysts and undeveloped fusomes (Fig 3B). One copy of *bam* was sufficient to rescue the *bam* null tumorous phenotype (Fig 3B).

Therefore the *D. ananassae* female *bam* null phenotype is consistent with that of *D. melanogaster, D. simulans*, and *D. yakuba*. In contrast, in *D. ananassae* males of the *bam* null genotype we did not observe over-proliferating spermatogonia (Fig 3B). The homozygous *bam* null genotype resulted in the same phenotype as the heterozygous bam null genotypes and wildtype indicating that *bam* is not necessary for the switch from proliferating spermatogonia to differentiated spermatocytes in *D. ananassae* males. Therefore, these cytological data are fully consistent with our fertility assay observations.

### Confirming the loss-of-function status of *bam* in the *bam* null disruption design

The only available Bam antibody is weakly cross-reactive in *D. melanogaster* and *D. simulans* thus we cannot directly demonstrate the absence of Bam protein in all the species tested (Flores 2013; Flores, Bubnell, et al. 2015). Although we used the same design to generate the *bam* null alleles across this study and therefore expect the 3xP3-DsRed and 3xP3-YFP disruptions to have the same effect on *bam* in all five species, we further confirmed the loss-of-function status of the *bam* disruption alleles in *D. ananassae* and *D. teissieri* given their functional divergence.

First, we asked if *bam* had undergone duplication events in *D. ananassae* and *D. teissieri* using publicly available sequence data and found no evidence of *bam* duplication in either species (details described in S1 File, S8 Table). Next, since the 3xP3-DsRed and 3xP3-YFP insertions introduce a premature termination codon, we expect the mRNA of the disrupted *bam* allele to be degraded through nonsense mediated mRNA decay (NMD), but if the premature termination codon induced exon skipping, the disrupted *bam* exon could be alternatively spliced out resulting in expression of a truncated *bam* allele, which may be sufficient for function (Brogna and Wen 2009; Mou et al. 2017; Chen et al. 2018; Sui et al. 2018). We used RT-qPCR to ask if patterns of expression in the *bam* null alleles in *D. ananassae* males and *D. teissieri* females and males were consistent with expectations for NMD. We found that for *D. ananassae* and *D. teissieri*, the expression of the *bam* null disruption alleles were reduced relative to the *bam* wildtype allele to levels consistent with NMD (∼1-50%, details in S1 File, Fig S1. Fig S2) (Pereverzev et al. 2015). Therefore, the mRNA of the *bam* disruption alleles are likely targeted for degradation. Additionally, since both *bam* null *D. teissieri* males and females show wildtype phenotypes we looked for evidence of exon skipping. We asked if the exon disrupted by 3xP3-YFP or 3xP3-DsRed in the *bam* null genotypes was expressed in proportion to the non-disrupted exons relative to the wildtype genotype. We found that the ratio of exon 1 to exon 2 expression was not significantly different between *D. teissieri* nulls and *D. teissieri* wildtype genotypes for males and females (details in S1 File, Fig S3).

Although *bam* null *D. ananassae* males and *D. teissieri* males and females exhibited wildtype phenotypes, together the genomic and RT-qPCR analyses indicate that the *bam* null alleles are loss-of-function alleles. This evidence therefore supports our conclusion that *bam* is not required for spermatogenesis in *D. ananassae* and *D. teissieri* nor oogenesis in *D. teissieri*.

### *D. ananassae bam* null males show evidence of germline defects

In *D. melanogaster*, Fas3 expression in the testes is limited to the hub, the testes microenvironment that houses the somatic and germline stem cell populations at the apical tip of the testes. As we found that *bam* was necessary for oogenesis in *D. ananassae* but not for spermatogenesis, we assayed for more subtle evidence of germline defects in *D. ananassae* males. We immunostained testes with an antibody to Fas3 to look for evidence of ectopic expression of Fas3 as in Gonzalez et al. (Gonzalez et al. 2015). In *D. melanogaster bam* null males, Fas3 is ectopically expressed, as the spermatogonia over proliferate (Fig 4A&B). In *D. ananassae bam* null males, we also observed ectopic expression of Fas3 in cells outside of the apical tip of the testes, although not nearly as severe as in *D. melanogaster bam* null males (Fig 4D&E). We then asked if there was any evidence of ectopic Fas3 expression in *D. teissieri bam* null males, as we also found that *bam* was not necessary for spermatogenesis in *D. teissieri*. In contrast to *D. ananassae* and *D. melanogaster*, we did not observe ectopic expression of Fas3 in *D. teissieri bam* null males (Fig 4B&C). This indicates that while *bam* is not necessary for differentiation in *D. ananassae* males, it may be playing a role in regulating the early differentiating germ cell population.

**Fig 4.**
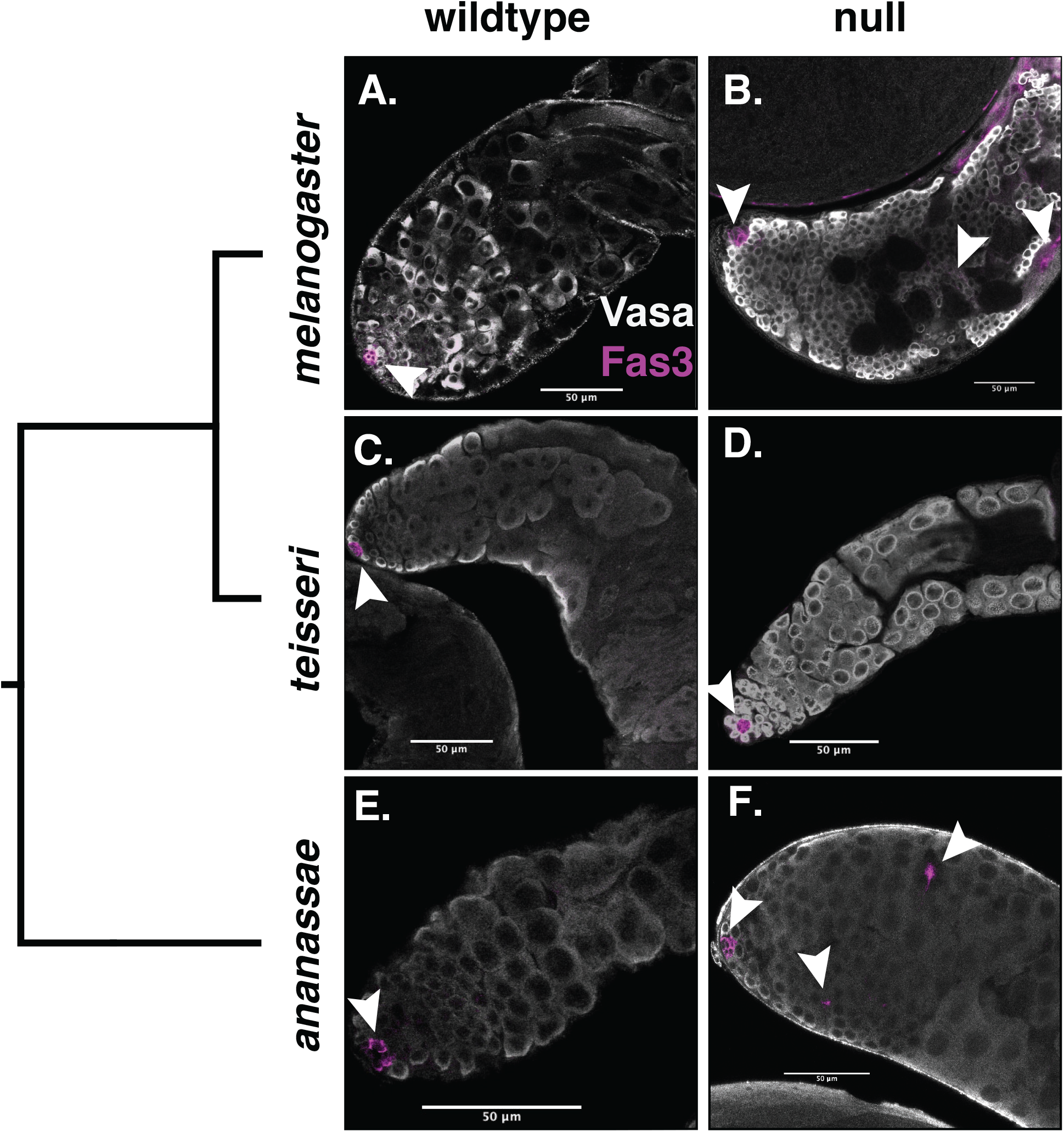
Loss of *bam* function in *D. ananassae* males shows ectopic expression of Fas3, a marker expressed in the hub, but not in *D. teissieri*. All testes tissue is labeled with antibodies for Vasa (germline) and Fas3 (hub, arrowheads). Significance was assessed with a Fisher’s exact test comparing counts of testes with ectopic Fas3 expression between wildtype and *bam* null testes. (A) *D. melanogaster bam* wildtype testes with Fas3 expression limited to the anterior tip of the testes hub (inset) and featuring large differentiating spermatogonia (Fas3 ectopic expression n=0/13). (B) *D. melanogaster bam* null testes feature ectopic Fas3 expression and small over-proliferating spermatogonia (Fas3 ectopic expression n=8/11, P<0.000). (C) *D. teissieri bam* wildtype testes with Fas3 expression limited to the anterior tip of the testes (hub) and featuring large differentiating spermatogonia (Fas3 ectopic expression n=0/20). (D) *D. teissieri bam* null testes with Fas3 expression limited to the anterior tip of the testes (hub) and large differentiating spermatogonia (Fas3 ectopic expression n=0/16, P=1). (E) *D. ananassae bam* wildtype testes with Fas3 expression limited to the hub and featuring large differentiating spermatogonia (Fas3 ectopic expression n=31). (F) *D. ananassae bam* null testes with ectopic Fas3 expression and large differentiating spermatogonia (Fas3 ectopic expression n=19/27, P<0.000).

### *bam* shows lineage-specific bursts of positive selection throughout the *Drosophila* genus

Previous studies have reported that *bam* is evolving under strong positive selection detected by the McDonald-Kreitman test in *D. melanogaster* and *D. simulans*, but not in *D. ananassae* (Civetta et al. 2006; Bauer DuMont et al. 2007; Choi and Aquadro 2014). As these three species span about 30 million years of divergence, we wanted to know if there was evidence of positive selection in lineages more closely related to *D. melanogaster* and *D. simulans*, or if this signal was unique to the individual *D. melanogaster* and *D. simulans* lineages. We expanded the species sampled to include three closely related species from the *yakuba* complex: *D. yakuba, D. santomea*, and *D. teissieri*. We also took advantage of published population genomic data from samples of inbred lines to expand our assessment of *bam* variation in additional population samples for *D. melanogaster* (Lack et al. 2015) and *D. simulans* and to newly assess *bam* variation in population samples in the *montium* species group (*D. serrata, D. bunnanda, D. birchii*, and *D. jambulina*) and the *ananassae* species subgroup (*D. bipectinata, D. pseudoananassae*, and *D. pandora*) (Li et al. 2021).

In addition to further sampling closely related species to *D. melanogaster* and *D. simulans*, we also wanted to ask if there was any evidence of adaptive evolution at *bam* in lineages outside of the *melanogaster* group. We resequenced *bam* from population samples of three more distantly related *Drosophila* species: *D. pseudoobscura, D. affinis*, and *D. mojavensis* which span approximately 70 million years of divergence (Li et al. 2021). We also used published population genomic data from samples of inbred lines to assess *bam* variation in *D. immigrans* and *D. rubida*, both members of the *immigrans* species group (Li et al. 2021).

To ask if any population samples showed departures from selective neutrality consistent with positive selection at *bam* similar to what we previously observed for *D. melanogaster* and *D. simulans*, we used a lineage-specific McDonald-Kreitman test (MKT) (Siddiq et al. 2017). The null hypothesis of the MKT is that the ratio of nonsynonymous to synonymous polymorphism within a species is equal to the ratio of nonsynonymous to synonymous fixed differences between species (or in our case, a predicted common ancestral sequence). A previous study reported signatures of positive selection at *bam* as an excess of nonsynonymous fixations in the *D. melanogaster* and *D. simulans* lineages (Bauer DuMont et al. 2007). As *bam* appears to be experiencing episodic signals of adaptive evolution restricted to individual lineages, we measured lineage-specific divergence. We used codeml to generate reconstructed ancestral sequences at the node of interest by maximum likelihood (Yang 1997; Yang 2007). We then used this sequence to calculate nonsynonymous and synonymous divergences for each population sample. We excluded polymorphisms below 12% as low-frequency mutations are likely to be slightly deleterious and reduce the power of the MKT to identify positive selection (Fay et al. 2002; Charlesworth and Eyre-Walker 2008).

For *D. melanogaster*, we utilized the Zambia population sample from the *Drosophila* genome Nexus to measure *bam* variation (Lack et al. 2015). The Zambia sample contains the largest sample size of intact *bam* sequence (few missing calls), and represents variation found in the ancestral range of *D. melanogaster*. We used the predicted common ancestor of *D. melanogaster* and *D. simulans* and found similar patterns of polymorphism and divergence to the smaller samples of *D. melanogaster* from the USA and Zimbabwe previously reported by Bauer DuMont et al. (Bauer DuMont et al. 2007) (Fig 5). The *D. melanogaster* lineage shows a strong signal of positive selection (alpha = 0.943, P=0.001) (Fig 5).

**Fig 5.**
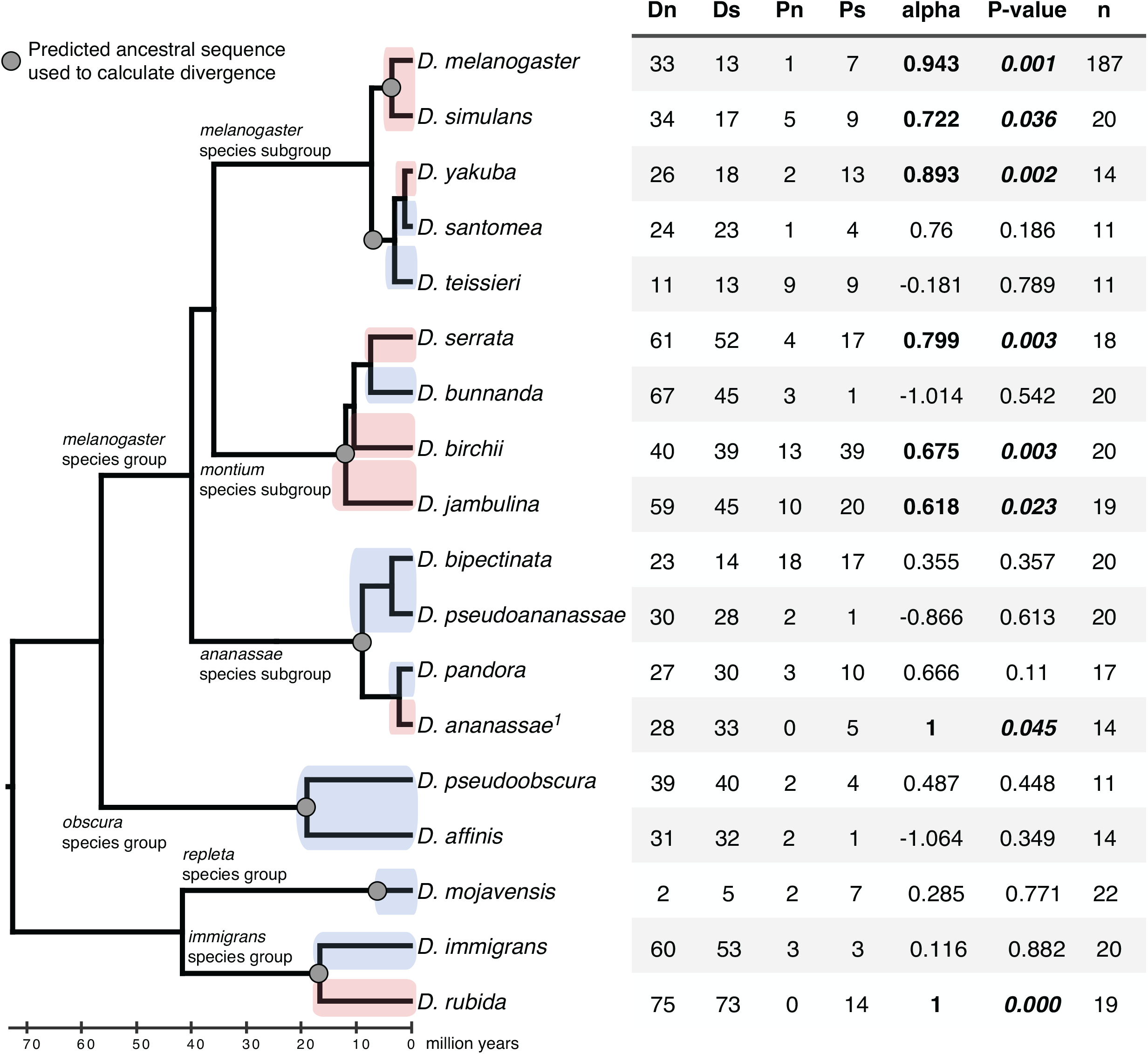
*Bam* shows lineage-specific signals of positive selection scattered across the *Drosophila* genus. Cladogram of *Drosophila* species we sampled in this study. Pink highlighted lineages represent lineages where we have evidence of positive selection at *bam* with the McDonald Kreitman test. Blue highlighted lineages are lineages of which we could not reject neutrality with the McDonald Kreitman test. The 2×2 table from the McDonald Kreitman test is shown on the right (N = nonsynonymous, S = synonymous). Alpha is the proportion of nonsynonymous fixations predicted to be due to positive selection. P-value is from the Chi-square test, n is sample size. 1Polymorphism data from Choi and Aquadro 2014.

For *D. simulans bam*, we utilized the Burnley, Victoria population sample from Li et al. (Li et al. 2021). We used the same common ancestral sequence as we did for the *D. melanogaster* analysis to calculate divergence. The *D. simulans* lineage shows a signal of positive selection for *bam* (alpha=0.722, P=0.036) are consistent with those previously reported by Bauer DuMont et al. (Bauer DuMont et al. 2007) (Fig 5). We also assessed *bam* variation in the *D. simulans* CA, USA population sample from Signor et al. (Signor et al. 2018), but due to elevated levels of genome-wide intermediate frequency polymorphisms caused by a recent population bottleneck, this sample violates key assumptions for the McDonald-Kreitman test (Fay 2011; Messer and Petrov 2013). For completeness, we report our findings for the CA sample in Supplementary File S1.

We used the predicted common ancestor of the *yakuba* complex (*D. yakuba, D. santomea, D. teissieri*) to estimate divergence for *bam* in each of the population samples. For *D. yakuba* we also observed a significant signal of positive selection for *bam* (alpha=0.893, P=0.002) (Fig 5). In contrast, we could not reject neutrality for *D. teissieri bam* or for *D. santomea bam*. The estimate of alpha is very low for *D. teissieri* (alpha=-0.011, P=0.983), but for *D. santomea* the alpha value was high, although not significant (alpha=0.76, P=0.186), which suggests it may be informative to sample *bam* from additional *D. santomea* populations with larger sample sizes (n=11 here) (Fig 5).

In the *montium* species subgroup, we used the predicted common ancestor of *D. serrata, D. bunnanda, D. birchii*, and *D. jambulina* to calculate divergence for *bam*. We found a significant signal of positive selection for *bam* in *D. serrata* (alpha=0.799, P=0.003), *D. birchii* (alpha=0.675, P=0.003), and *D. jambulina* (alpha=0.618, P=0.023) (Fig 5). However, we could not reject neutrality for *D. bunnanda* (alpha=-1.014, P=0.542) (Fig 5).

In the *ananassae* species group, we used the predicted common ancestor of *D. bipectinata, D. pseudoananassae, D. pandora*, and *D. ananassae* to calculate divergence for *bam*. We reassessed lineage-specific divergence at *bam* for the *D. ananassae* population sample previously reported by Choi et al (Choi and Aquadro 2014) as they reported divergence between *D. ananassae* and *D. atripex*. We did not reject neutrality at *bam* for *D. bipectinata* (alpha=0.355, P=0.357), *D. pseudoananassae* (alpha=-0.866, P=0.613), or *D. pandora* (alpha=0.666, P=0.11) (Fig 5). However, we found a significant signal of positive selection for *bam* in *D. ananassae* (alpha=1, P=0.045) (Fig 5). Therefore, the signals of positive selection at *bam* previously reported for *D. melanogaster* and *D. simulans* are not unique to the *melanogaster* species group.

In the *obscura* species group, we calculated divergence for *bam* from population samples of *D. pseudoobscura* and *D. affinis* from their predicted common ancestor and observed similar patterns of polymorphism and divergence (*D. pseudoobscura*: alpha=0.487, P=0.448), (*D. affinis*: alpha=-1.064, P=0.349), neither of which were close to rejecting neutral expectations (Fig 5). We used the predicted common ancestor of *D. mojavensis* and *D. arizonae bam* to calculate divergence in the *D. mojavensis* lineage and again did not observe a trend towards an excess of nonsynonymous divergence (alpha=0.285, P=0.771) (Fig 5). For the *immigrans* species group, we calculated divergence for *bam* from the predicted common ancestor of *D. immigrans* and *D. rubida*, We did not reject neutrality for *bam* in *D. immigrans* (alpha=0.116, P=0.882), but we did find a strong signal of positive selection for *bam* in *D. rubida* (alpha=1, P<0.000) (Fig 5). Therefore, the signals of positive selection at *bam* are heterogeneously distributed throughout the *Drosophila* genus.

## Discussion

### Bam’s role in germline stem cell maintenance and differentiation shows remarkable variation in the *Drosophila melanogaster* species group

Our comparative assessment of *bam* requirement in gametogenesis across diverse *Drosophila* species indicates that the functional evolution of *bam* is unexpectedly complex. We discovered that *D. teissieri* and *D. ananassae* do not require *bam* for gametogenesis in one or both sexes. A single orthologous copy of *bam* is found in all *Drosophila* species examined to date, but is absent in outgroups for example, house fly (Musca) and the sheep blow fly (Lucinia). These results raise the question as to whether *bam*’s critical role in GSC differentiation originally characterized in *D. melanogaster* is in fact a phylogenetically recent gain of function, or whether *bam*’s critical role is basal to the genus and that *bam*’s role has been recently (and possibly repeatedly) lost in specific lineages. For example, assuming *bam*’s role in GSC differentiation is basal, then the most parsimonious explanation of our functional data is that *bam* was required for oogenesis and spermatogenesis in the common ancestor of the species tested (Fig 1A), then *bam* function was lost in *D. ananassae* males, and then lost again independently in *D. teissieri* males and females. These alternative scenarios can be distinguished by future follow-up analyses of *bam* function in more divergent *Drosophila* species.

### Evidence of positive selection for protein diversification at Bam is strikingly heterogeneous across divergent *Drosophila* lineages

We analyzed additional population samples of species mainly within the *melanogaster* species group and a few more distant lineages for nucleotide sequence variation at *bam*. We found new evidence of lineage-specific signals positive selection for accelerated protein sequence evolution at *bam* in three of the *melanogaster* subgroup species (*D. melanogaster, D. simulans*, and *D. yakuba*), three of the *montium* subgroup species (*D. serrata, D. birchii*, and *D. jambulina*), and one of the *ananassae* subgroup species (*D. ananassae*). Additionally, we found evidence of positive selection in the *D. rubida* lineage, indicating that the protein diversification at *bam* is repeatedly occurring in species spanning over 70 million years of divergence.

While we could not reject neutrality for *bam* for the remaining lineages, It is important to note as well that the MKT has limited power statistically when applied to single genes (Akashi 1999; Zhai et al. 2009) and thus a failure to reject selective neutrality does not prove that positive selection is not acting or has not acted on this gene.

### Are stem cell genes hotspots for adaptive evolution?

The results of past studies of GSC evolution (Bauer DuMont et al. 2007; Choi and Aquadro 2014; Choi and Aquadro 2015; Flores, DuMont, et al. 2015) and our population genetic and functional results in this study lead to the question of why GSC developmental functions are frequent targets of adaptation. It has been documented that selfish transposable elements are upregulated in the absence of piRNAs, implying that piRNAs act to protect the germline genome leading to an evolutionary conflict that may drive positive selection at piRNA genes (Aravin et al. 2007; Simkin et al. 2013). There is natural variation in the gene *Bruno* that provides tolerance to deleterious P-element expression in the genome (Kelleher et al. 2018). In a scRNA-seq study of male *D. melanogaster* testes, Witt et al. (Witt et al. 2019) report many fixed de novo genes expressed in GSCs and early spermatogonia (where *bam* is expressed) at similar levels to testis-specific genes. In contrast, segregating de novo genes show lower expression in GSCs and early spermatogonia implying that cell type expression may impact the likelihood for a de novo gene to become fixed. As de novo gene evolution is increasingly recognized as a critical mode of adaptation, especially in reproduction, GSCs/early spermatogonia may be generating necessary variation for adaptation (Witt et al. 2019). A recent finding that the insect gene *oskar*, which is critical for germ cell formation and oogenesis, arose *de novo* through a horizontal gene transfer event from a prokaryote, resulting in a fusion between prokaryotic and eukaryotic sequence illustrates yet another mechanism by which novel genes can take on key developmental roles in reproduction (Jenny et al. 2006; Blondel et al. 2020).

The *bam* sequence itself has many properties of novel genes: it does not contain any known DNA or RNA binding motifs, it binds with multiple protein partners, contains structurally disordered regions, and is expressed in a tight spatiotemporal manner (Chen et al. 2010; Jiang and Assis 2017; Wilson et al. 2017; Klasberg et al. 2018; Vakirlis and McLysaght 2019). Novel genes that become fixed often gain critical roles in conserved gene networks. Their new role may lead to selection to fine tune their critical role thereby providing additional functional flexibility for these conserved processes (Chen et al. 2010). Such a process could represent what has been called developmental systems drift, a model to explain how developmental systems remain phenotypically conserved while the genetic pathways that determine them diverge (True and Haag 2001).

It is possible that *bam* was a gene novel to the common ancestor of *Drosophila* (and possibly some other dipterans) and has functionally diversified across the *Drosophila* genus, only gaining its role in germline stem cell differentiation in the lineage leading to the *melanogaster* species group. Additionally, it is possible that *bam* quickly evolved a role in GSC daughter differentiation in *Drosophila*, but that function has been shaped by germline conflicts in specific lineages. Conflict with germline agents (such as *W. pipientis)* may even drive the fixation of novel genes in new roles that successfully evade their control. As *W. pipientis* must recognize host cues during oogenesis to successfully infect the developing oocyte, it is possible that novel host genes that regulate differentiation might then be under selection for their new function if it evades unfavorable manipulation by *W. pipientis*.

It certainly seems plausible that genetic conflict is one way for new genes to evolve required functions in conserved processes such as GSC daughter differentiation. The *D. melanogaster* testes and the accessory gland show enriched expression of novel genes, many of which show signals of adaptive evolution, and some of which are predicted to be driven by sexual conflict (Levine et al. 2006; Zhou et al. 2008; Findlay et al. 2009; Kaessmann 2010; Gubala et al. 2017; Witt et al. 2019). While the developmental processes of gametogenesis and reproduction must be conserved, our results demonstrate that the individual genes that function in that process may vary among even closely related species.

## Materials and Methods

### Fly stocks and rearing

We raised fly stocks on standard cornmeal molasses food at room temperature. We used yeast glucose food for fertility assays. The *D. melanogaster w*^*1118*^ isogenic line and *w*^*1118*^ balancer lines were gifts from Luis Teixeira, and the *D. simulans w*^*501*^ line (stock 14021-0251.011), the *D. yakuba* white eyed line (14021-226.03) and the *D. ananassae* reference genome (14024-0371.14) lines are from the Drosophila species stock center <u>(h</u>t<u>tp://blogs.cornell.edu/drosophila/).</u>The *W. pipientis*-free *D. ananassae* line Nou83 was a gift from Artyom Kopp. The *D. teissieri* line (*teissieri* syn) was a gift from Daniel Matute (Turissini et al. 2015).

We tested all lines for *W. pipientis* infection status by qPCR (NEB Luna Universal qPCR kit) using primer sets for *wsp* and *arm* (S1 Table). Lines that tested positive (*D. teissieri, D. ananassae, D. simulans*) were treated with tetracycline (200 μg/mL) for three generations to remove *W. pipientis* (Glover et al. 1990). We allowed these lines to recover for at least three generations before using them in any analyses. We were unable to fully clear the *D. ananassae* reference line of *W. pipientis* with tetracycline, so to obtain a *W. pipientis* free stock, we crossed males of our *bam* null lines to females from the Nou83 line, which we confirmed to be *W. pipientis*-free by qPCR.

### Cloning for bam null constructs

We used Flybase to obtain the nucleotide sequence information for designing constructs in *D. melanogaster* (Dmel 5.57), *D. simulans* (dsim r2.01), *D. yakuba* (dyak r1.04), and *D. ananassae* (dana r1.04) (Thurmond et al. 2019). We obtained sequence information for the *D. teissieri* sequence from Turissini et al. 2017 (accession SAMN07407364-SAMN07407376) (Turissini and Matute 2017). We used Geneious for all cloning design.

We used the NEB Q5 High Fidelity 2X master mix to generate all PCR products. We gel extracted and purified PCR products using the NEB Monarch DNA gel extraction kit or Qiagen MinElute gel extraction kit. IDT primers were used for PCR, sequencing, and cloning. We generated the donor plasmids for the 3xP3-DsRed and 3xP3-YFP bam disruption lines using the NEB HiFi Assembly Cloning kit into the pHD-attP-DsRed vector from flyCRISPR (Gratz et al. 2014) as follows: we amplified 1.5 kb homology arms from genomic DNA of the appropriate species stock flanking the insertion site for 3xP3-DsRed or 3xP3-YFP. 3xP3-DsRed was amplified from the pHD-attP-DsRed plasmid and 3xP3-YFP was amplified from the *D. simulans* nos-Cas9 line, a gift from David Stern. We then gel extracted the two homology arms and the appropriate 3xP3 marker, purified them, and assembled them into the pHD vector backbone using the manufacturer’s protocol (S3 Table).

We used the U6:3 plasmid from CRISPRflydesign (Port et al. 2014) to express gRNAs for generating the *D. yakuba* lines. The gRNA target sequence was generated by annealing primers with the gRNA sequence and BbsI site overhangs, and ligating into the BbsI ligated U6:3 vector (T4 ligase NEB, BbsI NEB) (S3 Table). We used NEBalpha competent cells for transformations.

We prepped and purified all plasmids for embryo injections with the Qiagen plasmid plus midi-prep kit. We additionally purified plasmids for injections with phenol-chloroform extraction to remove residual RNases. We confirmed plasmid sequences with Sanger sequencing (Cornell BRC Genomics Facility).

### CRISPR/Cas9

#### gRNA selection

We chose gRNAs using the flyCRISPR target finder (Gratz et al. 2014) for *D. melanogaster, D. simulans, D. yakuba*, and *D. ananassae*. For *D. teissieri*, we used one completely conserved gRNA from *D. yakuba*. We also chose an additional gRNA from *D. yakuba* that had two divergent sites between *D. yakuba* and *D. teissieri*, thus we modified them to the *D. teissieri* sites. We selected gRNAs that had no predicted off-targets in the reference genome (S4 Table & S5 Table). For *D. yakuba*, we used the U6:3 gRNA plasmid from CRISPRfly design as described above. For the remaining species (*D. melanogaster, D. simulans, D. teissieri*, and *D. ananassae*), we used synthetic gRNAs (sgRNAs) from Synthego. We used 1-3 gRNAs per injection to increase the chances of a successful CRISPR event. We used multiple gRNAs if there were suitable options for that site to increase efficiency (multiple gRNAs within 50 bps with 0 predicted off-targets).

#### Injections

All CRISPR/Cas9 injections were carried out by Genetivision. Lines were injected with the plasmid donor, gRNAs (synthetic or plasmid), Cas9 protein (Synthego), and an siRNA for Lig4 (IDT DNA). The same siRNA was used for *D. melanogaster, D. simulans, D. yakuba*, and *D. teissieri* (pers communication with Daniel Barbash and David Stern S6 Table). We created a *D. ananassae*-specific siRNA due to sequence divergence (S6 Table).

For the *bam* disruption lines, we screened for eye color in F1s in-house using a Nightsea fluorescence system with YFP (cyan) and DsRed (green) filters. We backcrossed all CRISPR/Cas9 mutants to the stock line we used for injections for three generations. All mutants are maintained as heterozygous stocks. We confirmed all CRISPR insertions by Sanger sequencing (Cornell BRC Genomics Core Facility).

#### Genotypes for generating bam disruption homozygotes

We generated two *bam* disruption lines in each species in order to create the genotypes for the comparative *bam* null analyses. We generated one by disrupting *bam* with 3xP3-DsRed and the other with 3xP3-YFP. We selected DsRed or YFP positive flies and crossed them to generate homozygous *bam* disruption flies (*bam*^*3xp3-DsRed*^/*bam*^*3xP3-YFP*^). This cross also results in DsRed or YFP positive heterozygous *bam* null progeny as well as *bam* wildtype progeny without any fluorescent eye marker.

We targeted the 1^st^ exon of *bam* in all species in order to disrupt the *bam* coding sequence and introduce an early stop codon. While we were successful in generating 1^st^ exon disruptions in each species, in *D. simulans* we were only able to obtain a 3xP3-YFP 1^st^ exon disruption line. However, we did find that a *D. simulans* line with a 3xP3-DsRed insertion in its 2^nd^ intron resulted in homozygous sterile females and males with tumorous ovaries and testes, while heterozygous females and males were fertile with wildtype ovary and testes phenotypes. As this edit was specific to the *bam* locus, we presumed it was an mRNA null allele and we used this line for our analyses. Thus, if the phenotype of the 3xP3-DsRed 2^nd^ intron allele over the 3xP3-YFP 1^st^ exon allele is the same as the homozygous 3xP3-DsRed 2^nd^ intron allele, this is consistent with classification as a genetic null.

### Fertility assays

#### Female fertility

All female fertility assays were performed following the approach of Flores et al. (Flores et al. 2015). Specifically, we collected virgin females and aged them based on their time to sexual maturity 2-3 days (*D. melanogaster, D. simulans, D. yakuba*, and *D. teissieri*) or 5-7 days (*D. ananassae*) (Markow and O’Grady 2006). We collected all genotypes from each bottle to control for bottle effects. We collected virgin males of the wildtype genotype for each species and aged them for the same timeframe as the females of that species. We evenly distributed males collected from different bottles across the female genotypes to control for any bottle effects. We crossed single females collected within 48 hours of each other to two virgin males. We allowed the trio to mate for 9 days, and then flipped them to new vials. Except for *D. ananassae* and *D. yakuba*, as fertility was very low, we cleared the trio after day 9 and did not continue further. For all other species crosses, after an additional 9 days, we cleared the trio. Then, we counted the progeny for each trio every two to three days to get the total adult progeny per female.

#### Male fertility

All male fertility assays were performed as described for the female fertility assays except that one tester male was crossed to two virgin females.

#### Food and conditions for fertility assays

For *D. melanogaster, D. simulans, D. ananassae*, and *D. teissieri*, we performed all fertility assays on yeast-glucose food. For *D. yakuba*, we performed fertility assays on cornmeal-molasses food, as this line did not thrive on the yeast-glucose food. We kept all crosses and fertility experiments in an incubator at 25°C with a 12-hour light-dark cycle.

### Statistics

For the fertility assays we used estimation statistics to assess the mean difference (effect size) of adult progeny between the wildtype *bam* genotype and each individual additional *bam* genotype. For generating the estimation statistics and plots (Ho et al. 2019), we used the dabest package in Python (v. 0.3.1) with 5,000 bootstrap resamples. Estimation statistics provide a non-parametric alternative to other statistical tests of the difference of the mean (for example, ANOVA), and allow us to understand the magnitude of the effect of *bam* genotype on the fertility phenotype. In text, we report significance as a mean difference (effect size) outside the 95% bootstrap confidence interval. While one benefit of these statistics is that they are less biased than reporting traditional P-values, we also report P-values for approximate permutation tests with 5,000 permutations.

### Immunostaining

We used the following primary antibodies, anti-vasa antibody from Santa Cruz biologicals (anti rabbit 1:200), anti-Hts-1B1 from Developmental Studies Hybridoma bank (anti-mouse 1:4 for serum, 1:40 for concentrate item), and anti-Fas3 from DSHB (anti-mouse 1:50). We used the following secondary antibodies: Alexaflour 488, 568 (Invitrogen) at 1:500.

We performed immunostaining as described in Aruna et al. and Flores et al. (Aruna et al. 2009; Flores, Bubnell, et al. 2015). Briefly, we dissected ovaries and testes in ice-cold 1X PBS and pipetted ovaries up and down to improve antibody permeability. We fixed tissues in 4% paraformaldehyde, washed with PBST, blocked in PBTA (Alfa Aesar, with 0.1% Triton X added), and then incubated in the appropriate primary antibody in PBTA overnight. We then washed (PBST), blocked (PBTA), and incubated the tissue in the appropriate secondary antibody for 2 hours, then washed (PBST) and mounted in mounting media with DAPI (Vecta shield or Prolong Glass with NucBlue) for imaging.

### Confocal Microscopy

We imaged ovaries and testes on a Zeiss i880 confocal microscope with 405nm, 488 nm, and 568 nm laser lines at 40X (Plan-Apochromat 1.4 NA, oil) (Cornell BRC Imaging Core Facility). We analyzed and edited images using Fiji (ImageJ).

### RNA extraction

For RT-qPCR experiments we dissected ovaries from 10 one-day old females per biological replicate or testes from 15 one-day old males in ice cold 1x PBS. The tissue was immediately placed in NEB DNA/RNA stabilization buffer, homogenized with a bead homogenizer, and then stored at -80°C until RNA extraction. We used the NEB Total RNA mini-prep kit for all RNA extractions according to the manufacturer’s protocol and included the on column DNAse treatment. We assessed RNA concentration and quality with a Qubit 4 fluorometer and a NanoDrop.

### RT-qPCR

For all RT-qPCR analyses, we used the Luna One Step RT-qPCR kit from NEB according to the manufacturer’s instructions at 10μl total volume in 384 well plates. We used a Viia7 qPCR machine with the appropriate cycle settings as described in the Luna One Step RT-qPCR kit.

For the analysis of *bam* expression in *D. ananassae*, we used primers to *D. ananassae bam* with *D. ananassae Rp49* as a control (S1 Table). To assess the possibility that the *bam* disruption allele induced exon skipping of the disrupted exon 1 in *D. teissieri*, we designed a primer pair that spanned exon 1 to exon 2 and another pair that spanned exon 2 to exon 3 (S1 Table). We used the exon 2 - exon 3 target as the control. For both the *D. teissieri* and *D. ananassae* assays described above, we included a standard curve for each target on every plate at a 1:2 dilution and then used the standard curve method to calculate the relative quantity of *bam* (to either *Rp49* for *D. ananassae* or *bam* exon 2-exon 3 for *D. teissieri*) with the QuantStudio software using the wildtype *bam* genotype as a control for each species.

For the allele specific analysis in *D. teissieri*, we designed one primer pair specific to the *bam* 3xP3-DsRed and 3xP3-YFP insertion alleles and another primer pair specific to the wildtype allele. We then used these two targets to measure the relative expression of the insertion allele to the wildtype allele in a single heterozygous sample. The primer pair for the *bam* insertion alleles target the sequence between the SV40-polyA tail of the 3xP3-DsRed and 3xP3-YFP insertion and the *bam* sequence downstream of the insertion (S1 Table). The primer pair for the *bam* wildtype allele target *bam* upstream and downstream of the 3xP3-DsRed and 3xP3-YFP insertion, and therefore will not amplify the large insertion allele under RT-qPCR conditions (S1 Table). We included a standard curve for each target on every plate at a 1:2 dilution and then used the standard curve method in the QuantStudio software to calculate the Ct and Ct standard error for each target and each sample. We report the relative quantity of the 3xP3-DsRed or 3xP3-YFP allele using the wildtype allele in the same sample as a control by calculating 2^-(nullCt-wildtypeCt)^.

### Genomic analysis of *bam* copy number in *D. teissieri*

While *bam* is reported as a 1:1 syntenic ortholog on FlyBase for *D. melanogaster, D. simulans, D. yakuba*, and *D. ananassae*, data for *D. teissieri* is not available. To ask if *bam* has been recently duplicated in *D. teissieri*, we generated a blast database (blastn v 2.9.0) from a new de novo assembled *D. teissieri* reference genome (accession GCA_016746235.1) and blasted *D. teissieri bam* from the resequencing described above.

Additionally, to assess read coverage for *bam* compared to the surrounding loci and the entire chromosome 3R, we used publicly available short read WGS data for *D. teissieri* (SRR13202235) and used bwa-mem (v 0.7.17) to map reads to a new de novo assembled *D. yakuba* reference genome (accession GCA_016746365.1). We visualized the alignment and assessed read coverage using Geneious.

### Population genetic analyses

The *D. yakuba, D. santomea*, and *D. teissieri* population sample lines were gifted by Brandon Cooper and Daniel Matute (Cooper et al. 2017). The *D. yakuba* and *D. teissieri* lines were collected in Bioko in 2013, the *D. santomea* sample was collected in Sao Tome in 2015. The *D. affinis* population sample lines were gifted by Robert Unckless (S2 Table). The *D. mojavensis* population sample is from Organ Pipe National Monument, Arizona and DNA from this sample was gifted by Erin Kelleher and Luciano Matzkin. The *D. pseudoobscura* population sample is from Apple Hill, California, as described in (Schaeffer and Miller 1991). We used previously prepared DNA from the *D. pseudoobscura* population sample as described in (Schaeffer and Miller 1991).

We prepared genomic DNA from *D. affinis, D. yakuba, D. santomea*, and *D. teissieri* samples using the Qiagen PureGene kit. We used Primer3 to generate primers to amplify *bam* from *D. affinis, D. mojavensis, D. pseudoobscura, D. yakuba, D. santomea*, and *D. teissieri* population samples (S7 Table). We used existing polymorphism data (Daniel Matute, personal communication) to generate primers for *D. yakuba, D. santomea*, and *D. teissieri*. We resequenced *bam* using Sanger sequencing from 5’UTR-3’UTR for *D. yakuba, D. santomea*, and *D. teissieri* and *bam* exons and introns for *D. affinis* (Cornell BRC Genomics Facility). We used Geneious for aligning the sequences and calling polymorphisms. We generated fasta files for each individual. We used DnaSP to phase individuals with heterozygous sites, and then randomly picked one allele for further analyses (Bauer DuMont et al. 2007). We will deposit all resequencing data described here in Genbank.

We used polymorphism data for *D. melanogaster* from the Zambia population sample from the Drosophila genome Nexus (Lack et al. 2015), as this is the largest population sample with intact *bam* sequence and represents a population in the *D. melanogaster* ancestral range. For the *D. melanogaster* Zambia population sample, we removed a single individual that contained missing nucleotide calls in the *bam* coding sequence (ZI117). As we filtered polymorphisms at < 12% frequency, omitting this single individual would not affect our downstream analyses.

Polymorphism for *D. simulans, D. serrata, D. bunnanda, D. birchii, D. jambulina, D. bipectinata, D. pseudoananassae, D. pandora, D. immigrans*, and *D. rubida* as well as the single *bam* sequence for *Z. bogoriensis* that we used as an outgroup are from Li et al 2021(accession PRJNA736147) (Li et al. 2021). *D. ananassae bam* polymorphism, and single *D. atripex bam* sequences are from Choi et al. (Choi and Aquadro 2014). We obtained single *bam* sequences for species used for outgroups from FlyBase (*D. erecta*), and Genbank (*D. eugracilis, D. arizonae, D. navojoa, D. miranda, D. subobscura)*.

We performed all multiple sequence alignments for the *bam* coding sequences using PRANK (v.170427) with the -codon and -F parameters and using the PRANK generated guide tree. As *bam* is highly divergent across the *Drosophila* genus, we generated multiple sequence alignments only among closely related clusters of species to ensure alignment quality. We generated a multiple sequence alignment of the *melanogaster* subgroup species (*D. melanogaster, D. simulans, D. yakuba, D. santomea, D. teissieri, D. erecta, D. eugracilis*) and *montium* subgroup species (*D. serrata, D. bunnanda, D. birchii, D. jambulina*), a multiple sequence alignment of *D. ananassae, D. bipectinata, D. pseudoananassae, D. pandora*, and *D. atripex*, a multiple sequence alignment of *D. mojavensis, D. arizonae, and D. navajo*, a multiple sequence alignment of *D. affinis, D. pseudoobscura, D. miranda*, and *D. subobscura*, and a multiple sequence alignment of *D*.*immigrans, D. rubida*, and *Z. bogoriensis*.

We used only unique sequences for input to codeml. We used the codeml package from PAML (v. 4.9) (Yang 1997; Yang 2007) to generate the predicted common ancestor sequences to calculate lineage-specific divergence for *bam* with the McDonald-Kreitman test. We used the PRANK alignments and their corresponding gene trees as input to codeml with the following control file parameters (noisy = 9, verbose = 2, runmode = 0, seqtype = 1, CodonFreq = 2, clock = 0, aaDist = 0, model = 0, NSsites = 0, icode = 0, getSE = 0, RateAncestor = 1, Small_Diff = .5e-6, cleandata = 0, method = 1).

We used the http://mkt.uab.cat/mkt/mkt.asp webtool to perform the standard McDonald-Kreitman test (MKT) comparing nonsynonymous vs. synonymous changes (Egea et al. 2008). We excluded polymorphic sites at < 12% frequency, as these are likely slightly deleterious alleles that have not yet been purged by purifying selection (Charlesworth and Eyre-Walker 2008). For each species where we had polymorphism data, we performed the MKT using the specified predicted common ancestral sequence to calculate lineage-specific divergence. We report the values of the contingency table, the P-value of the Chi-Square test, as well as alpha, the proportion of fixations predicted to be due to positive selection (Eyre-Walker 2006).

### Summary of accession numbers for genomes used in this analysis

For read coverage analysis of *bam*: *D. teissieri*: SRR13202235, GCA_016746235.1. *D. yakuba*: GCA_016746365.1. For cloning *D. teissieri bam* null constructs: SAMN07407364-SAMN07407376. For *bam* polymorphism from *D. simulans, D. serrata, D. bunnanda, D. birchii, D. jambulina, D. bipectinata, D. pseudoananassae, D. pandora, D. immigrans, and D. rubida*: PRJNA736147.

## Acknowledgements

We thank Santiago Herrera Alvarez for resequencing *bam* from *D. affinis* and Jae Young Choi for resequencing *bam* from *D. pseudoobscura* and *D. mojavensis*. We thank Daniel Matute, Brandon Cooper, Robert Unckless, Erin Kelleher, and Luciano Matzkin for the population samples they provided for this study. We thank Fang Li for working to make the variant call files from (Li et al. 2021) quickly available for this study. Additionally, we thank David L. Stern for his advice and guidance in designing the *bam* null alleles and for an extremely thoughtful and critical review of an earlier draft of the manuscript. We are grateful to Mariana F. Wolfner, Andrew G. Clark, Miwa Wenzel and Catherine Kagemann for valuable input and discussion of these experiments and results and helpful comments on this manuscript. Lastly, we thank the three anonymous reviewers for their thoughtful and insightful feedback for improving this manuscript.

## Data Availability

Sequence data generated in this study will be deposited in GenBank prior to publication. The remaining data underlying this article are available in the article and in its online supplementary material. Fly lines, CRISPR/Cas9 reagents, and plasmids used in this study are available upon request.

---

**Fig S1.**
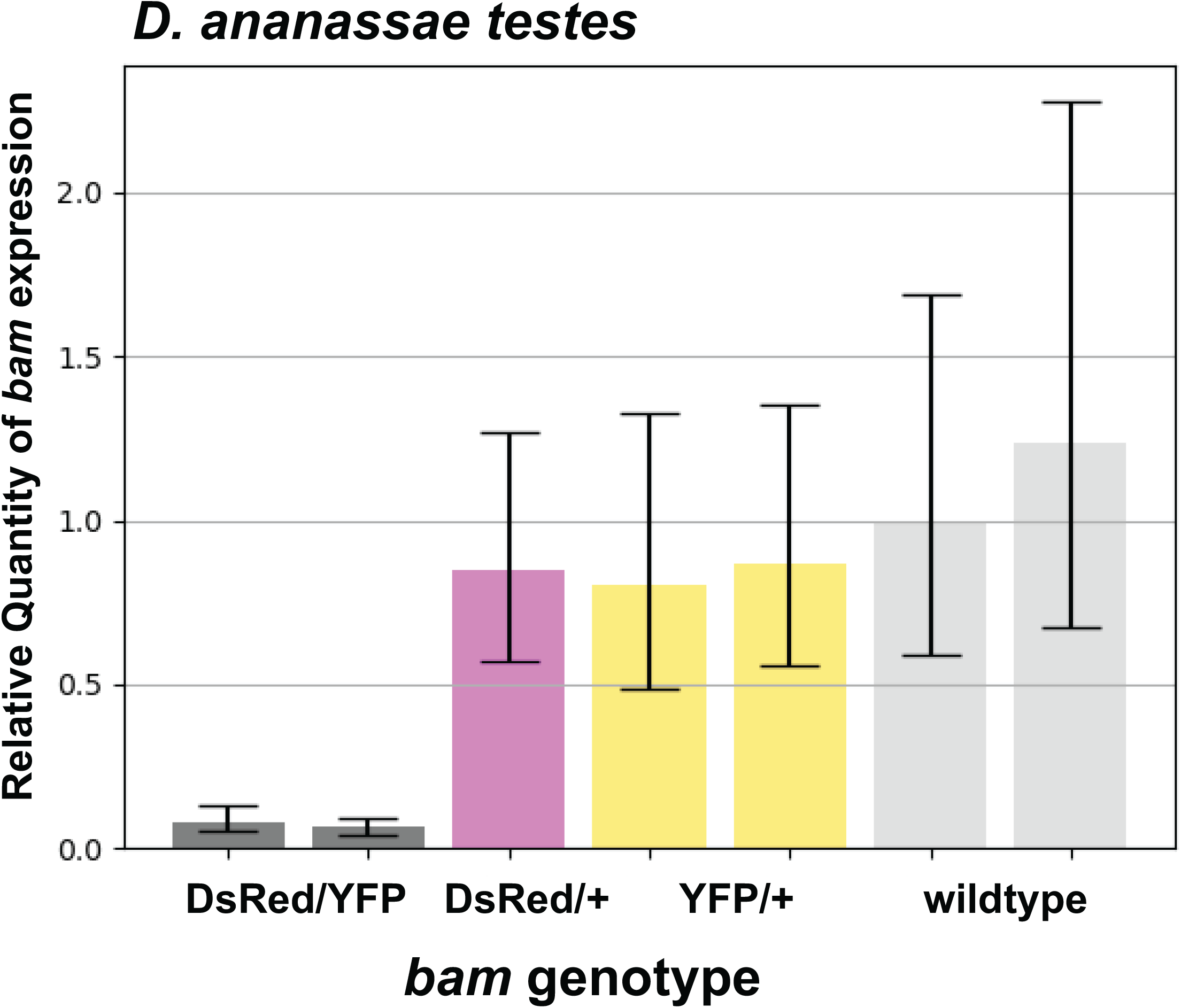
RT-qPCR experiment to assess *bam* mRNA in *D. ananassae bam* null genotypes reported as the relative quantity (RQ) of *bam* expression to the housekeeping control RP49. *bam* expression in testes of the null (DsRed/YFP) genotype is significantly reduced (standard error) relative to *bam* expression in testes of the wildtype genotype at approximately 10%, consistent with expectations for NMD. The heterozygous genotypes DsRed/+ and YFP/+ do not show reduced *bam* expression relative to wildtype.

**Fig S2.**
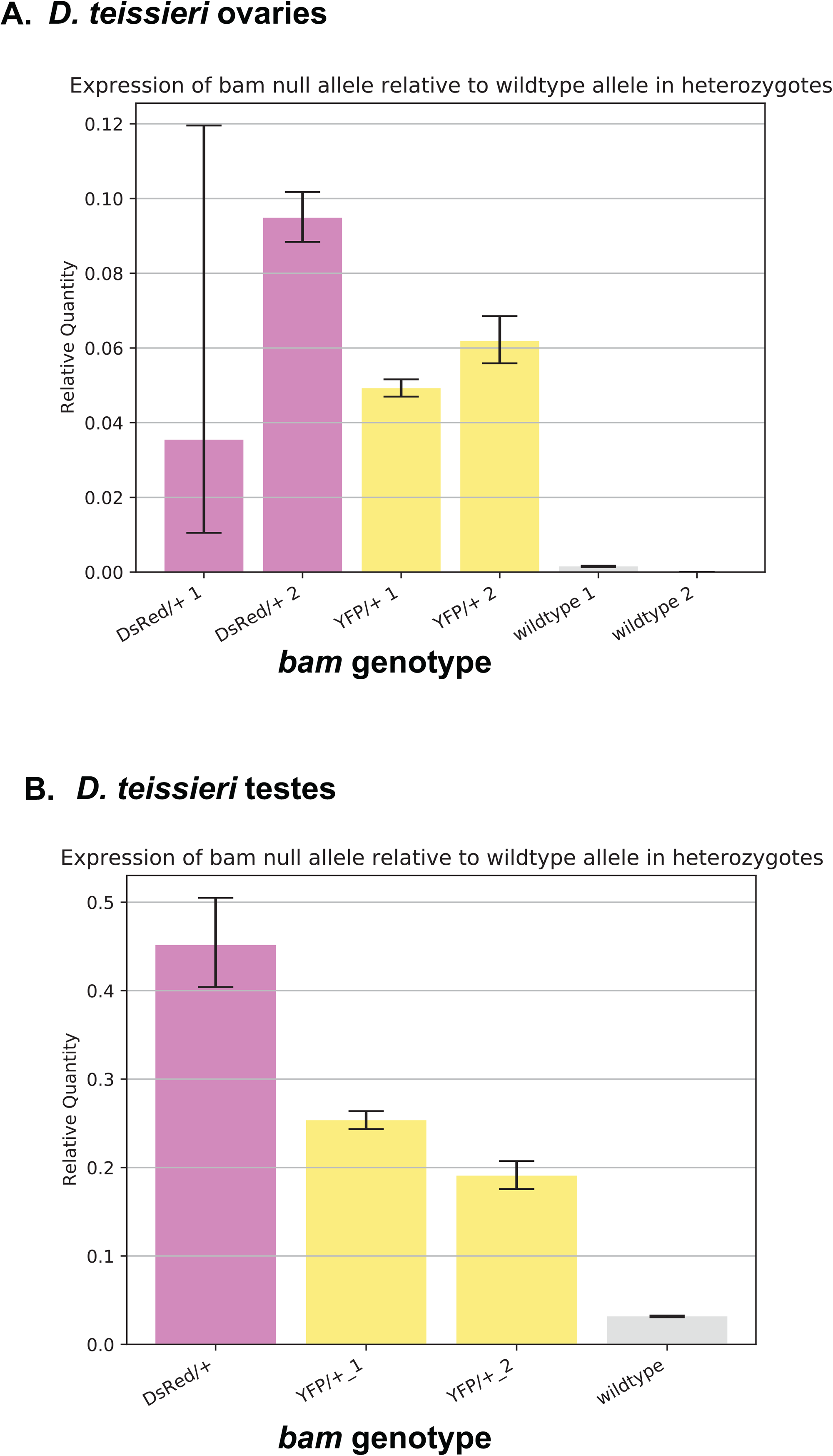
RT-qPCR experiment to detect patterns consistent with nonsense mediated mRNA decay of the *bam* disruption allele in *D. teissieri* reported as the relative quantity (RQ) of mRNA from the *bam* disruption allele to the *bam* wildtype allele in heterozygotes. Significance reported as standard error (A) Expression of the *bam* null allele relative to the *bam* wildtype allele in ovaries. Both heterozygous genotypes DsRed/+ and YFP/+ show less than 10% of expression of the *bam* null allele relative to the wildtype allele, consistent with expectations for NMD. The wildtype negative control shows little to no expression of the null allele as expected. (B) Expression of the *bam* null allele relative to the *bam* wildtype allele in testes. Both heterozygous genotypes DsRed/+ and YFP/+ show less than 50% of expression of the *bam* null allele relative to the wildtype allele, consistent with expectations for NMD. The wildtype negative control shows little to no expression of the null allele as expected.

**Fig S3.**
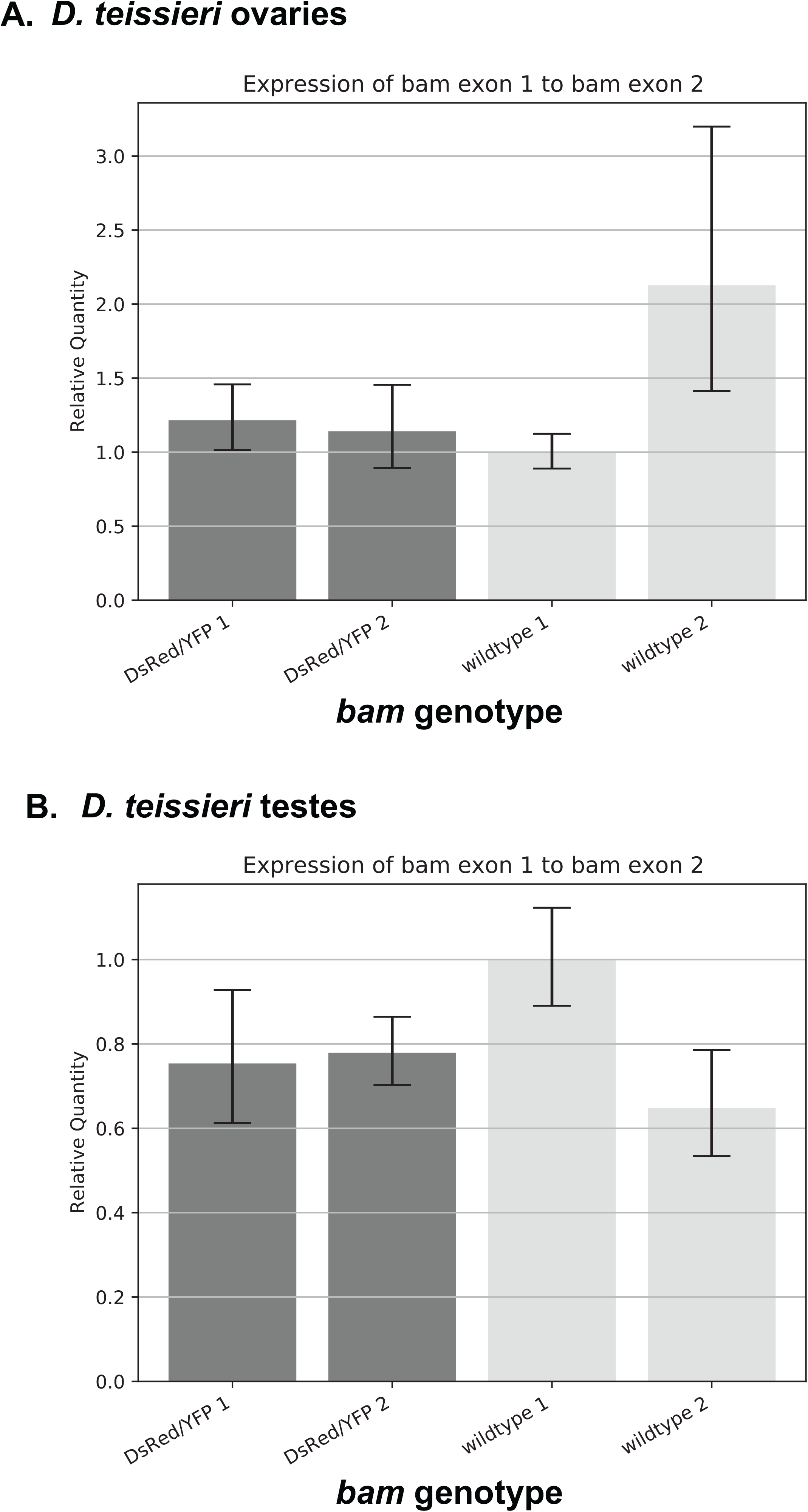
RT-qPCR experiment to detect evidence of exon skipping of the exon containing the *bam* disruption reported as the relative quantity (RQ) of *bam* exon 1 to exon 2. Significance reported as standard error (A) Expression of *bam* exon 1 relative to exon 2 in ovaries. The RQ of exon 1 to exon 2 in the *bam* null DsRed/YFP genotype is not significantly different than the RQ of exon 1 to exon 2 in the *bam* wildtype genotype, indicating that expression of the exon containing the disruption is comparable to the unaffected exon in both the null and wildtype genotype, and therefore exon skipping is not likely occurring in females. (B) Expression of *bam* exon 1 relative to exon 2 in testes. The RQ of exon 1 to exon 2 in the *bam* null DsRed/YFP genotype is not significantly different than the RQ of exon 1 to exon 2 in the *bam* wildtype genotype, indicating that expression of the exon containing the disruption is comparable to the unaffected exon in both the null and wildtype genotype, and therefore exon skipping is not likely occurring in males.

