## Supplementary File S1 for "Functional divergence of the *bag of marbles* gene in the *Drosophila melanogaster* species group"

**Supplementary File 1**

**Contents:**

**Confirming the loss of function status of *bam....................................................*2**

**Non-equilibrium population history for *D. simulans* CA, USA...........................5**

**Confirming the loss-of-function status of *bam* in the *bam* null disruption design:**

**No evidence for *bam* duplications in *D. ananassae* and *D. teissieri* genomes:**

If *bam* underwent a duplication in *D. ananassae* or *D. teissieri* and we only disrupted one *bam* copy, that could explain our observations that *bam* is not necessary for GSC daughter differentiation in these species, as there could be another functional copy of *bam*. Notably, FlyBase reports *D. ananassae bam* is a 1:1 syntenic ortholog (along with *D. melanogaster, D. simulans,* and *D. yakuba bam*). Data for *D. teissieri* is not reported in FlyBase, so we used a new *D. teissieri* de novo reference genome (accession in Methods) to create a blast database and assess sequence similarity to *bam* across the genome. We used the *bam* resequencing data for *D. teissieri* described above to blast the *D. teissieri* genome and found only one location mapping to *bam* on chromosome 3R, syntenic with *bam* from the other *Drosophila* species reported (data not shown). While the reference genome was generated with long reads (Pac Bio) and thus even a recent duplication should be assembled, we wanted to ensure that finding *bam* in single copy in *D. teissieri* was not a result of mis-assembly. We used publicly available deep sequencing short read data from the same *D. teissieri* line and mapped it to a new *D. yakuba* de novo reference genome (accession in Methods) and asked if read coverage was elevated around *bam* compared to the surrounding loci and the entire chromosome arm. We found that *bam* and its surrounding genomic region is within the mean read coverage for chromosome 3R, further indicating that *bam* is single copy in the *D. teissieri* genome (S8 Table).

**Expression levels of *bam* null disruption alleles in *D. ananassae* males, *D. teissieri* females, and *D. teissieri* males is consistent with expectations for nonsense mediated mRNA decay:**

Additionally, if our *bam* disruption design did not cause a *bam* loss of function allele in *D. ananassae* and *D. teissieri,* that could also explain our results. To generate our *bam* null alleles we used CRISPR/Cas9 to insert a 3xP3-YFP or 3xP3-DsRed gene cassette into the first exon (except for the 3xP3-DsRed *D. simulans* line, which is in the second intron). We chose this method over deleting the entire coding sequence because it had been described in the literature (Gratz et al. 2014) as a suitable method to generate null alleles, and due to concerns that large deletions can disrupt the regulation of neighboring genes. Notably, *bam* shares a 3’UTR with its neighboring gene. We expect that the early termination codon introduced by the 3xP3 promoter sequence (and more in the subsequent gene cassette) will trigger nonsense mediated mRNA decay (NMD), and thereby a loss of function allele (Brogna and Wen 2009). However, as the only existing Bam antibody is weak in *D. melanogaster* and *D. simulans*, we cannot confirm the absence of Bam protein in all of the species tested. Additionally, a variety of studies have reported that the introduction of premature termination codons by genome editing can induce the conserved cellular process of exon-skipping (Mou et al. 2017). This has been reported in a variety of species (human, mouse, rabbit, locust) although not yet in *Drosophila* species (Mou et al. 2017; Chen et al. 2018; Sui et al. 2018). In some exons, premature termination sequences result in alternative splicing and excision of the affected exon. If the next exon is in-frame, or there is a suitable site for translation initiation, it is possible to express a truncated version of the protein (Mou et al. 2017). This phenomenon has not been reported for edits that result in a premature termination codon in exon 1, but regardless we want to rule out this event as an explanation for our observations in *D. teissieri* and *D. ananassae*. Additionally, if this phenomenon occurred and was generating a truncated Bam protein, it may not be recognized by the weak Bam antibody (polyclonal), so the absence of immunostaining would not rule out exon skipping.

We first hypothesized that if the null allele was targeted by NMD, mRNA from the *bam* null alleles should be reduced at approximately 1-50% compared to mRNA from wildtype *bam* alleles (Pereverzev et al. 2015). We tested this in *D. ananassae* males where we performed RT-qPCR for *bam* on testes for all genotypes reported here (wildtype, DsRed/+, YFP/+, DsRed/YFP) using Rp49 as a housekeeping control. We used the standard curve method to measure the relative quantity of *bam* expression using the wildtype genotype as the control. We found that *bam* expression in null testes is approximately 10% of *bam* expression in wildtype and heterozygous genotypes (Fig S2). In *D. ananassae bam* null testes, we do not see over-proliferation of spermatogonia despite our result that *bam* expression in the null males is reduced to 10%. This indicates that the disruption design likely results in the degradation of *bam* mRNA in testes.

Due to inconsistencies in optimizing primers for *D. teissieri* Rp49 as a housekeeping gene, and to design an assay that would control for the number of cells expressing *bam* for future studies (as *bam* null and wildtype tissue differ in cell types), we designed an assay to assess NMD that would not be affected by differences in cell types or tissue morphology and does not rely on a housekeeping gene to quantify expression. We used the heterozygous genotypes, which across all species tested here exhibit wildtype tissue morphology, to ask if the *bam* disrupted allele was degraded at levels consistent with NMD (1-50%) relative to the wildtype allele. We used the standard curve method and calculated the ∆Ct between the *bam* null allele and *bam* wildtype allele in each sample as a measure of relative quantity. We performed this allele specific RT-qPCR for *bam* in *D. teissieri bam* null heterozygotes and a wildtype control and found that in both DsRed/+ and YFP/+ heterozygous ovaries expression of the *bam* null allele is less than 20% of the wildtype *bam* allele (Fig S3 A), and therefore consistent with expectations for NMD. In DsRed/+ and YFP/+ heterozygous testes, we found that expression of the *bam* null allele is less than 50% of the wildtype *bam* allele, and therefore also consistent with expectations for NMD (Fig S3 B). These results indicate that the *bam* disruption allele is likely targeted for degradation at the mRNA level.

**No evidence for exon skipping in *D. teissieri* females and males:**

Since the presence of NMD does not completely rule out exon skipping, and both *bam* null *D. teissieri* females and males showed wildtype phenotypes, we additionally asked if the relative expression of *bam* exon 1 to *bam* exon 2 was consistent between the *D. teissieri bam* null genotype and the *D. teissieri bam* wildtype genotype. If exon skipping is not occurring, we would expect the relative expression of exon 1 to exon 2 in the *bam* null genotype to be approximately 1:1 with the relative expression of exon 1 to exon 2 in the wildtype *bam* genotype. We used RT-qPCR to amplify sequence that spanned exon 1 - exon 2 and exon 2 - exon 3 in *bam* null ovaries and testes and wildtype *bam* ovaries and testes. We found that in *bam* null ovaries, the expression of exon 1 to exon 2 is not significantly different from wildtype expression levels (Fig S4 A). Therefore, it is unlikely that exon skipping of the disrupted exon is occurring in females. In *bam* null testes, we also found no significant difference in the expression of exon 1 to exon 2 compared to wildtype expression (Fig S4 B). Therefore, it is also unlikely that exon skipping of the disrupted exon is occurring in males. Taken with the allele specific assay, the *bam* disruption allele in *D. teissieri* is likely being degraded without exon skipping. Therefore, it is unlikely that the *bam* null genotype is expressing any functional *bam* gene product in *D. teissieri.*

**Non-equilibrium population history for the *D. simulans* CA population may affect the power of the McDonald-Kreitman test**

For *D. simulans bam*, we utilized the CA, USA population sample from Signor et al (Signor et al. 2018). We previous reported significant departure from an equilibrium neutral model using McDonald-Kreitman test in the direction consistent with positive selection for amino acid sequence diversification at *bam* for a sample of size n = 10 chromosomes from *D. simulans* from North Carolina (Aquadro et al. 1988; Bauer DuMont et al. 2007). For the current study, we additionally analyzed two new population samples of *D. simulans*, one from California (Signor et al. 2018) and another from Australia recently published by Li et al. (2021). We used the same common ancestral sequence as we did for the *D. melanogaster* analysis to calculate divergence. However, we did not observe the same patterns of divergence and polymorphism in this sample as we previously observed in the NC sample In the California *bam* sample, we cannot reject neutrality (Dn=46, Ds=25, Pn=13, Ps=15; alpha = 0.539; P=0.084), in contrast to our previous finding in the NC sample (Dn=26, Ds=16, Pn=9, Ps=15; alpha=0.631; P=0.049). It is notable that although the ratios were not significantly different, the trend in this lineage also shows a high alpha value and higher nonsynonymous divergences. The difference between the two samples seems to be driven by higher nonsynonymous polymorphism in the California sample compared to the NC sample.

The failure to reject selective neutrality for *bam* in the *D. simulans* population sample from California appears to be due to a recent non-equilibrium population history for the the species in California. There is a genome-wide excess of intermediate frequency polymorphisms in the California population sample that is consistent with a combination of population contraction and possibly soft sweeps on standing genetic variation within the past 100 years in this California *D. simulans* population (Signor et al. 2018). The introduction of the *w*Ri strain of *W. pipientis* into southern California around the 1980s led to the spread of this strain through CA with initially strong cytoplasmic incompatibility (Hoffmann et al. 1986) that would have resulted in a severe bottleneck particularly in the female effective population size (e.g. (Cariou et al. 2017)) and perhaps accounts for the strong departure from equilibrium in this sampled California population. A population bottleneck can bring slightly deleterious polymorphisms to higher frequency. While the frequency of these deleterious polymorphisms will then slowly decline, during this time their frequency in the population is thus an overestimation of their probability of their fixation and thus weaken power of the MKT and underestimate alpha (e.g. (Messer and Petrov 2013)). For this reason, we report in this manuscript the results from the sample of size n=20 chromosomes sequenced from the single Australian population (Burnley, Victoria) by Li et al. (2021).
